# Environmental DNA Surveys of Invertebrate Community on Forest Canopies Using Rainwater Analysis

**DOI:** 10.1101/2024.08.11.607477

**Authors:** Takumaru Miwa, Naoya Miyashita, Chisato Numa, Hideyuki Doi

## Abstract

Forest canopies, known for their high biodiversity, are essential for understanding forest ecosystems. Traditional methods for canopy surveys, such as tree climbing and canopy walkways, face challenges related to safety, cost, and time constraints. Environmental DNA (eDNA) analysis, which involves examining DNA from environmental samples, offers a promising alternative for these surveys. This study investigates the feasibility of using rainwater to collect eDNA from forest canopies, utilizing rain’s natural ability to wash away DNA from hard-to-reach areas. By comparing DNA analysis results from rainwater with conventional records obtained through visual and capture surveys, this research aims to validate the effectiveness and reliability of this method. Preliminary findings suggest that eDNA analysis from rainwater could provide an efficient approach to canopy biodiversity surveys, though further validation is required. This study marks an important first step towards developing eDNA analysis as a complementary tool for forest canopy research.

## Introduction

Forest canopy is the top portion of the forest, the set of branches and foliage, and is a highly biodiverse area where a wide variety of animals live. Because of the localization of ecosystem functions, it is essential to understand the forest canopy biota to understand the biodiversity of forests (e.g., Basset et al. 2012, Lowman et al. 2012, Schowalter and Chao 2021). However, due to the difficulty of accessing the forest canopy for collection and other purposes, all kinds of methods have been introduced for biological surveys. Various access methods have been developed and introduced to the canopy, including tree climbing using rope techniques, large canopy access equipment such as canopy walkways (e.g., Lowman et al. 2012, Schowalter and Chao 2021), canopy cranes, and jungle gyms, as well as advanced technology such as small balloons and airships. These have made it possible to directly conduct tree canopy surveys following various research sites (e.g., Basset et al. 2012). However, these forest canopy surveys have problems such as safety considerations, financial costs, and survey time.

Environmental DNA (eDNA) is a generic term for DNA contained in soil and water, and the biological monitoring method to analyze this eDNA to evaluate the presence/absence and biomass of species is called eDNA analysis (e.g., Beng and Corlett 2023, Doi et al. 2021, Takahara et al. 2012, Takahara et al. 2013). To solve such issues and conduct efficient forest canopy biological surveys, this study attempted to conduct biological surveys of the forest canopy using eDNA analysis (Aucone et al. 2023, Ladin et al. 2021, Macher et al. 2022, Valentin et al. 2020). eDNA analysis can be performed by collecting water samples in the environment and examining the DNA information in them, which has several advantages in terms of cost and work efficiency compared to conventional survey methods such as sampling and visual inspection (e.g., Stemhagen et al. 2024). The eDNA analysis has been conducted mainly by collecting environmental water samples, but there have been some difficulties in introducing eDNA analysis to forest surveys, such as the limited water area and limited scope for surveys in forests (Ladin et al. 2021, Macher et al. 2022). Considering these problems, this study attempted to implement eDNA analysis in forest canopy biological surveys by collecting water from rain that falls on the forest canopy. Based on the case study of spraying water on shrubs to wash away eDNA attached to shrub leaves and collecting liquid containing eDNA (Valentin et al. 2020, Yoneya et al. 2023), rain, a natural product, may be able to wash away eDNA in the forest canopy, which is difficult for humans to reach (Ladin et al. 2021, Macher et al. 2022). In addition, a research paper on the use of drones to collect eDNA directly from branches in the forest canopy, which confirmed a case of less eDNA being collected when conducted after rainfall, discusses the possibility that rain has washed away the eDNA in the forest canopy (Ladin et al. 2021, Macher et al. 2022).

Rain falling on the forest canopy may wash away eDNA attached to tree trunks, leaves, and branches, and the collection of rainwater containing eDNA may allow for eDNA analysis of the forest community. Such studies of eDNA analysis from rainwater have been conducted in recent years (Ladin et al. 2021, Macher et al. 2022). This method has the potential to be more efficient than conventional forest canopy biological surveys because rain falls over a wider area, allowing surveys to be conducted over a wider area, and because it is not necessary to access the forest canopy, which has been a problem in forest canopy surveys. However, the method has not been sufficiently tested against traditional surveys such as visual and capturing surveys, highlighting the need for further validation to confirm its effectiveness and reliability.

The purpose of this study is to develop eDNA analysis as more efficient visual survey method for the forest canopy. Here, we compare the results of this study with the recent biological survey records. By evaluating the agreement or disagreement with the visual/capturing surveys for organisms detected in the eDNA survey, the potential applications of this study are clarified.

## Materials and methods

### Sampling Location

We conducted the water sampling in the forests surrounding the Himeji City Science Museum in Himeji City, Hyogo Prefecture, Japan (N 34.85173, E 134.62688). The precipitation data recorded at the water collection survey sites is presented in Table 1. The data was obtained from the Himeji Special Regional Meteorological Observatory (N 34.8395, E 134.6710) in Himeji City, Hyogo Prefecture (Japan Meteorological Agency 2023). The data was collected on December 21-22, 2022 (hereafter, Winter survey), and June 8-9, 2023 (Summer). The geodetic distance between the Himeji Special Regional Meteorological Observatory and the survey site at the Himeji City Science Museum (N 34.85173, E 134.62688) is 4.258 km.

**Table 1.**
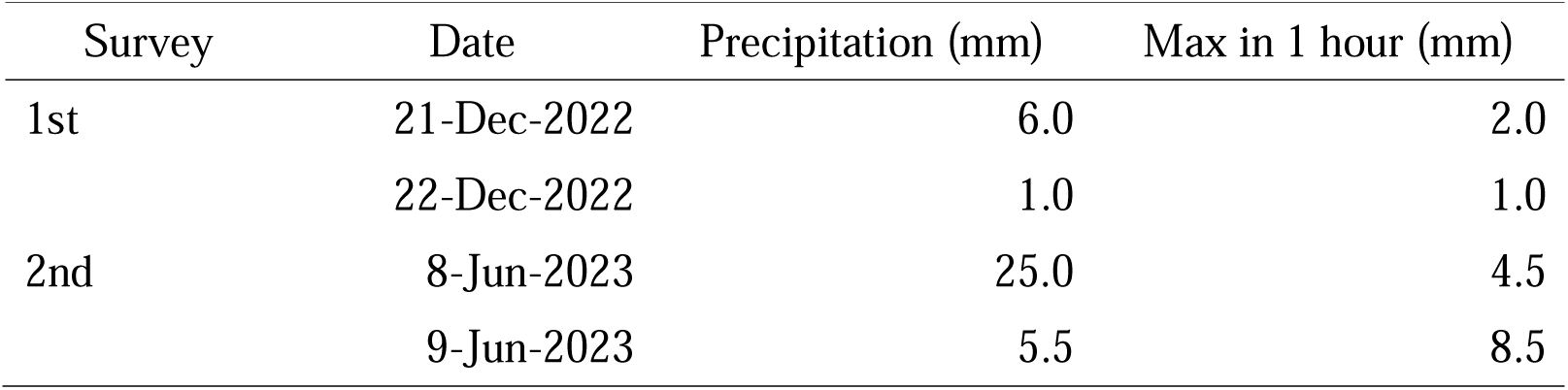
The precipitation data recorded at the water collection survey sites.

### Water sampling

Water sampling was conducted on December 21-22, 2022, and June 8-9, 2023, totaling two rounds of sampling. The sampling method involved setting up buckets before precipitation events, with the first round on December 21, 2022, and the second round on June 8, 2023. Buckets were placed under the canopy of ten evergreen trees to obtain ten samples of canopy-passing rainwater (Fig. 1). Additionally, four samples were collected from streams near the trees at the same locations, one sample of rainwater not passing through the canopy was collected in December, and two samples were collected near the Himeji City Science Museum in June. Field blanks were also collected to assess contamination resulting from sampling and transportation methods, where samples were obtained from locations devoid of the target organism’s DNA in the environment [9]. In this study, 100 mL of purified water was poured into 200 mL disposable plastic cups, and filtration was performed in a similar manner. The collection locations of each sample are shown on the map in Fig. 1. In the December survey, rainwater not passing through the canopy was collected under deciduous trees, while in the June survey, considering the influence of the dense canopy in spring, sampling locations were changed to near the buildings of the Himeji City Science Museum. To discuss the tree species through which the rain passed, the tree species near the bucket installations were examined and recorded (Fig. 1). Four locations were identified as belonging to the Fagaceae family, three to the Betulaceae family, and one each to the Machilaceae, Quercaceae, and Theaceae families.

**Figure 1.**
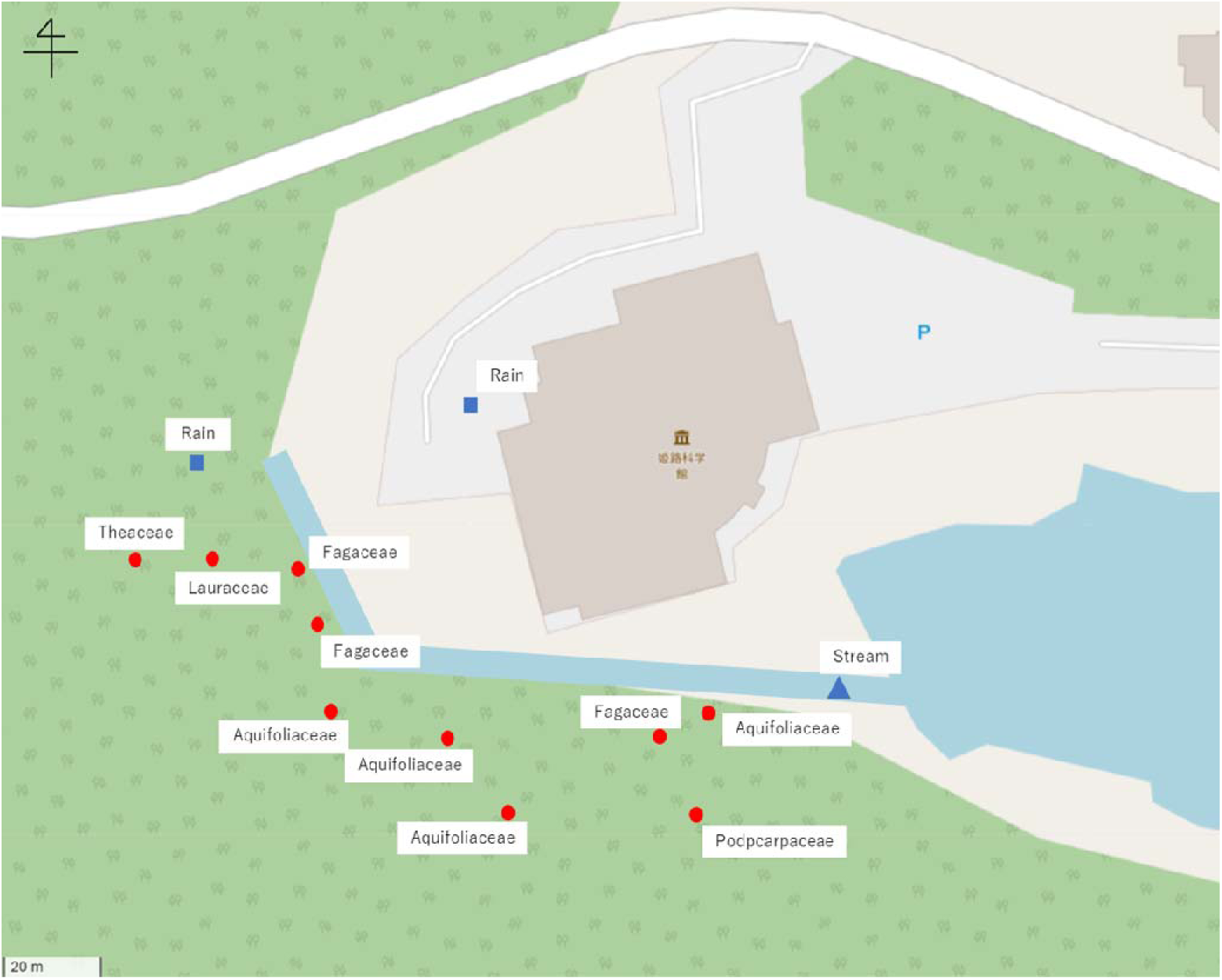
The map of collection locations. Red points represent the tree securing the buckets for water collection. Rain and Stream mean rain and stream water collections. The map was obtained by OpenStreetMap (https://www.openstreetmap.org).

The bucket installation method involved securing the buckets with pegs in locations conducive to collecting rainwater passing through the canopy (Fig. 2). When installing the buckets, it was considered that the entry of fallen leaves could lead to a higher proportion of eDNA derived from them, potentially affecting the analysis results. Indeed, in a preliminary survey conducted on November 30, 2022, before the December survey, it was found that the period until sufficient rain fell after bucket installation on the same day was long, allowing the entry of fallen leaves. Starting from the next survey, measures were considered to prevent the entry of fallen leaves, such as using nets, but it was recognized that this would introduce a problem where rainwater would always pass through fallen leaves if fallen leaves accumulated on the net. Therefore, bucket installations were continued without adding tools. However, for both the December and June surveys, buckets were installed and collected within 24 hours to prevent the entry of fallen leaves.

**Figure 2.**
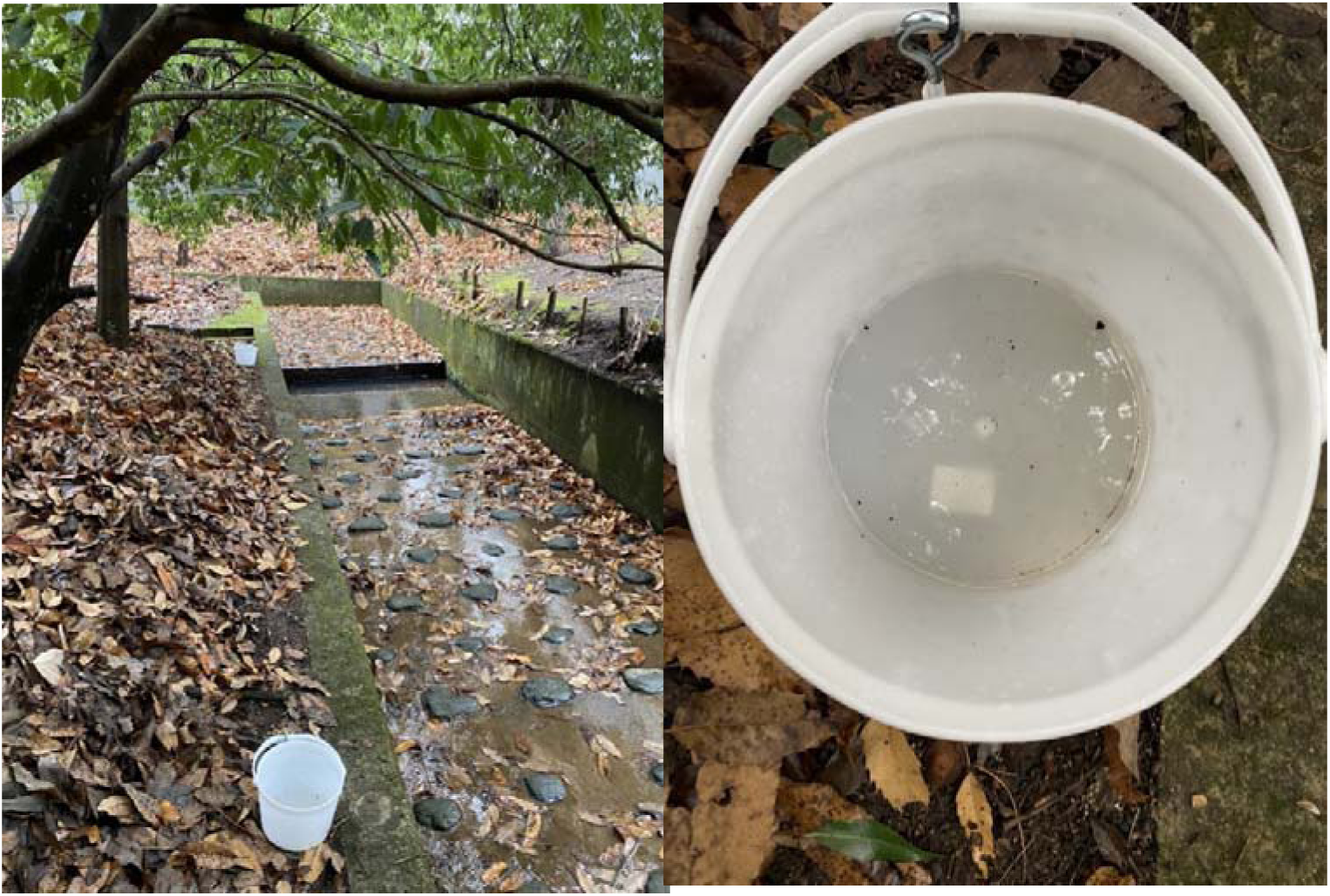
The photos of bucket installation.

### Water Sampling Method

The water collection and filtration process were initiated once a sufficient quantity of rainwater had accumulated in the buckets. The method for water collection and filtration from the accumulated rainwater in the buckets involved on-site filtration using Sterivex filters according to Minamoto et al. (2021). A 50 mL syringe was used to collect 50 mL of rainwater from the buckets. The collected rainwater was then injected from the 50 mL syringe into the injection port of the Sterivex filter (GV 0.22 µm, Millipore, Billerica, MA, USA) for filtration. This process was repeated twice, resulting in a total filtration volume of 100 mL. If a significant amount of material remained in the Sterivex filter after the first filtration, the second filtration required more time. Subsequently, efforts were made to remove as much moisture remaining inside the Sterivex filter as possible using the air pressure of the syringe. Once the moisture inside the Sterivex filter was nearly removed, the Sterivex filter was sealed onto the syringe, and its outlet port was sealed using a lure fitting. RNAlater (Sigma-Aldrich, St. Louis, MI, USA) was then added to the syringe, and using air pressure, RNAlater was injected into the Sterivex filter. Once the Sterivex filter was filled with RNAlater, the injection port was sealed using a lure fitting. For samples collected from streams, water collection was performed directly from the stream using a 50 mL syringe, followed by filtration of 100 mL. Sterivex filters subjected to these procedures were placed in a cooler box containing sufficient ice packs for preservation and transported back to the laboratory, where they were stored at –20°C.

### DNA Extraction

DNA extraction from Sterivex filters obtained on-site was performed using the DNeasy Blood and Tissue kit (Qiagen, Hilden, Germany) according to Miya et al. (2016). Initially, RNAlater within the Sterivex was extracted using a 50 mL syringe. Subsequently, a premix was prepared by combining 20 µL of Proteinase-K (600 mAU/mL), 200 µL of Buffer AL, and 220 µL of PBS(-) per Sterivex unit (totaling 440 µL), which was then injected into the Sterivex. The injected Sterivex units were incubated in a rotator within an oven, rotated at 10 rpm, and heated to 56°C for 20 minutes. Following this, the Sterivex inlet was inserted into a 2.0 mL tube, and each Sterivex with its corresponding tube was placed into a 50 mL conical tube. The conical tube was capped and centrifuged at 6,000 g for 5 minutes. Subsequently, 200 µL of ethanol was added to the 2.0 mL tube containing the extracted DNA, thoroughly mixed using a pipette, and then transferred into a column. The column was centrifuged at 6,000 g for 1 min. After centrifugation, the collection tube was replaced, Buffer AW1 was added, and centrifuged again at 6,000 g for 1 min. Following this, Buffer AW2 was added, and centrifuged at 20,000 g for 3 min. After centrifugation, the collection tube was discarded, and the column was placed into a 1.5 mL tube. Subsequently, 100 µL of elution Buffer AE was added onto the membrane of the column, incubated at room temperature for 1 minute, and centrifuged at 6,000 g for 1 min. This process was repeated twice to achieve a total elution volume of 200 µL. After centrifugation, the column was removed, and the 1.5 mL tube containing the eluted DNA was stored at –20°C.

### Metabarcoding Analysis

Metabarcoding analysis through high-throughput sequencing was performed using the Illumina MiSeq platform. The 1st PCR involved amplifying the target region using sequence primers and universal primers containing random bases N. The primer sequences are listed in Table 1. The premix for each sample consisted of 0.2 µL of KOD FX Neo (TOYOBO, Osaka, Japan), 6.0 µL of 2× PCR Buffer for KOD FX Neo (TOYOBO), 2.4 µL of 2 mM dNTPs (TOYOBO), and 0.7 µL each of Primer Forward and Reverse, totaling 10.0 µL. The premixes were mixed with 2.0 µL of thawed template DNA, followed by PCR using a Mastercycler nexus gradient (Eppendorf, Hamburg, Germany). Invertebrates, encompassing organisms devoid of a backbone or vertebrae, include the class Insecta within their classification. As this study focuses on invertebrates inhabiting the canopy, primers specifically targeting invertebrates, particularly those of the class Insecta, were selected. The primer (Forward: Fwh_F2 and Reverse: EPTD_r2n) with the highest proportion of invertebrates sequences among those described by Leese et al. (2021) was chosen. The primer sequence described in Leese et al. (2021) including random bases N at the 5’ terminus, were designed for the 1st PCR to target these primers. For the 1st PCR cycles, adjustments were made for each survey. For samples surveyed on December 21st to 22nd, 2022, the 1st PCR cycle comprised an initial denaturation at 95°C for 2 min, followed by 40 cycles of denaturation at 95°C for 30 sec, annealing at 46°C for 45 sec, and extension at 68°C for 45 sec. A final extension was conducted at 68°C for 5 min before cooling to 10°C. For samples surveyed on June 8th to 9th, 2023, adjustments were made to the cycles due to insufficient amplification. The 1st PCR cycle involved an initial denaturation at 94°C for 2 min, followed by 8 cycles of denaturation at 95°C for 30 sec, annealing at 60°C for 1 minute and 30 sec, and extension at 72°C for 2 min. This was followed by 24 cycles of denaturation at 95°C for 30 sec, annealing at 48°C for 1 minute and 30 sec, and extension at 72°C for 2 min. Subsequently, 23 cycles of denaturation at 95°C for 30 sec, annealing at 60°C for 1 min and 30 sec, and extension at 72°C for 2 min were performed. Finally, an extension at 68°C for 5 min was conducted before cooling to 10°C. The products of the 1st PCR were purified using AMPure XP (Beckman Coulter, Brea, CA, USA) and used as templates for the 2nd PCR. The 2nd PCR was conducted to attach MiSeq adapters and index sequences to the 1st PCR products. The premix for the 2nd PCR consisted of 6.0 µL of KAPA HiFi HS Ready Mix (Kapa Biosystems, Cape Town, South Africa) and 1.4 µL each of tagged 2nd PCR Primer Forward and Reverse, totaling 8.8 µL. These premixes were mixed with 3.2 µL of template DNA, followed by PCR using a Mastercycler nexus gradient (Eppendorf). The 2nd PCR cycle involved an initial denaturation at 95°C for 3 min, followed by 12 cycles of denaturation at 98°C for 20 sec, and extension at 72°C for 15 sec. A final extension was conducted at 72°C for 5 min before cooling to 10°C. The indexed 2nd PCR products were purified again using AMPure XP (Beckman Coulter, Brea, CA, USA), and DNA libraries were sequenced on the MiSeq platform using the MiSeq Reagent Kit v3 300 bp pair-end sequencing (Illumina, San Diego, CA, USA).

## Data Analysis

Demultiplexed fastq files for the sequence data were processed using DADA2 ver. 1.26.0 package (Callahan et al. 2016) in R ver. 4.2.0 (R Core Team 2022) and Claident (Tanabe and Toju 2013). DADA2 is capable of accurately distinguishing sequences from each sample, even with a difference of only one base pair. By utilizing DADA2, the misinterpretation of sequences generated by sequencing errors as biologically derived sequences can be minimized. Initially, quality filtering and trimming were performed to remove low-quality reads and adjust to specified trimming sizes. Subsequently, dereplication was conducted to eliminate duplicate sequences from the sequence data and extract unique sequences along with their counts. Following this, sample inference was performed using the quality scores of unique sequences to derive reliable sample information, and an error model was created. This error model helps minimize the impact of sequencing errors on the analysis results, thereby enhancing data reliability. Pair-end reads were merged to obtain more reliable sequence information.

Subsequently, the removal of chimeric sequences, which are unnatural sequences generated during processes such as PCR reactions, was conducted. DADA2’s ability to process each sequence individually enables the detection of related chimeras, facilitating more accurate chimera removal. Next, using the results obtained from DADA2, taxonomic assignment was performed using Claident. Claident maintains databases for taxonomic assignments at different classification hierarchies and resolutions, utilizing the “overall_class” database for identification.

## Observation Survey

We used the observation survey records from 2019 to 2023. Surveys were conducted mainly around the vicinity of the Himeji City Science Museum. The records were compiled using visual observations, photography, and specimen preservation. Survey methods included direct collection using sweep nets and hands as well as bait trapping using bananas, pupa powder or canned tuna. If capturing certain organisms proved challenging, visual observations or photography were used for recording. All specimens were preserved as dry specimens. Invertebrates’ specimens were subject to appropriate methods of euthanasia, including ethyl acetate or freezing for insects other than Lepidoptera, and thoracic compression for most Lepidopteran species.

## Statistical analysis

To demonstrate community similarity, we employed vegan ver. 2.6.4 package. We organized the data to represent the presence/absence information of taxa at each sampling location and time, thereby transforming it into binary data for each taxonomic group. We conducted non-metric multidimensional scaling (NMDS) using Jaccard distance with metaMDS function. We aimed for a Stress value less than 0.1, indicative of good fit, while a value greater than 0.2 would suggest poor fit. We selected the trial with the smallest Stress value among 20 randomizations for NMDS analysis. NMDS visually represents similar communities proximate and dissimilar communities distant in the plot. We plotted community similarity among samples for taxonomic groups in a two-dimensional space using the obtained coordinates. We evaluated statistical significance using PERmutational Multivariate ANalysis Of VAriance (PERMANOVA), examining how community similarity was influenced by multiple factors. Comparisons were made between the sequence read counts of taxonomic groups per tree species and the total sequence read counts of all detected taxonomic groups to calculate the abundance of taxonomic groups per tree species. Visualizations were achieved using the functions in ggplot2 version 3.4.4.

## Results

We detected total 1,248 taxa in all sequence samples. Although algae and other organisms were also detected in the analysis, this study primarily focuses on invertebrates species, hence they are not included in the results. Also, the detected taxonomy levels in invertebrates species were mostly family level or higher. So, we used family as the taxonomy level to show the results. The results of the Winter survey revealed detection of 19 different families, with Diptera, Coleoptera, Neuroptera, and Acari being the most frequently detected orders (Table 2). Samples from the canopy drip rain yielded a higher diversity of detected families, while no families were confirmed from the rain samples. Samples from the stream yielded six detected families, with all but Chironomidae and Corixidae being exclusive to stream samples. Chironomidae was notably prevalent in canopy drip rain and stream samples, showing significantly higher sequence read counts compared to others. Additionally, benthic organisms such as Cambaridae and Lumbricidae were detected even in canopy drip rain samples.

**Table 3.** The list of detected sequence reads of the invertebrate families in the different categorical samples in Winter survey.

| Family | Rain | Rain under open<br>canopy | Stream |
| --- | --- | --- | --- |
| Chironomidae | 102702 | 0 | 10054 |
| Mycetophilidae | 8933 | 0 | 0 |
| Caeciliusidae | 2794 | 0 | 0 |
| Tipulidae | 141 | 0 | 2325 |
| Bdellidae | 2203 | 0 | 0 |
| Ceratopogonidae | 1946 | 0 | 0 |
| Lepidostomatidae | 0 | 0 | 1862 |
| Simuliidae | 1486 | 0 | 0 |
| Apataniidae | 0 | 0 | 907 |
| Nemouridae | 0 | 0 | 752 |
| Ixodidae | 266 | 0 | 0 |
| Enchytraeidae | 248 | 0 | 0 |
| Megascolecidae | 0 | 0 | 151 |
| Nymphalidae | 66 | 0 | 0 |
| Chaoboridae | 52 | 0 | 0 |
| Cambaridae | 34 | 0 | 0 |
| Nepticulidae | 30 | 0 | 0 |
| Baetidae | 21 | 0 | 0 |
| Psychidae | 9 | 0 | 0 |

Summer survey results were similar to the winter survey. the results of the summer survey are presented in Table 3. Ten families were detected, with Diptera being the most prevalent. The summer survey also showed a higher prevalence of families detected from canopy drip rain samples, with Chironomidae being notably abundant, a family not detected in the winter survey. Additionally, Coleoptera, mainly inhabiting tree trunks and decaying wood, was detected. Rain samples predominantly yielded families such as Potamidae, while stream samples revealed Lumbricidae, with some families overlapping with canopy drip rain samples.

**Table 3.**
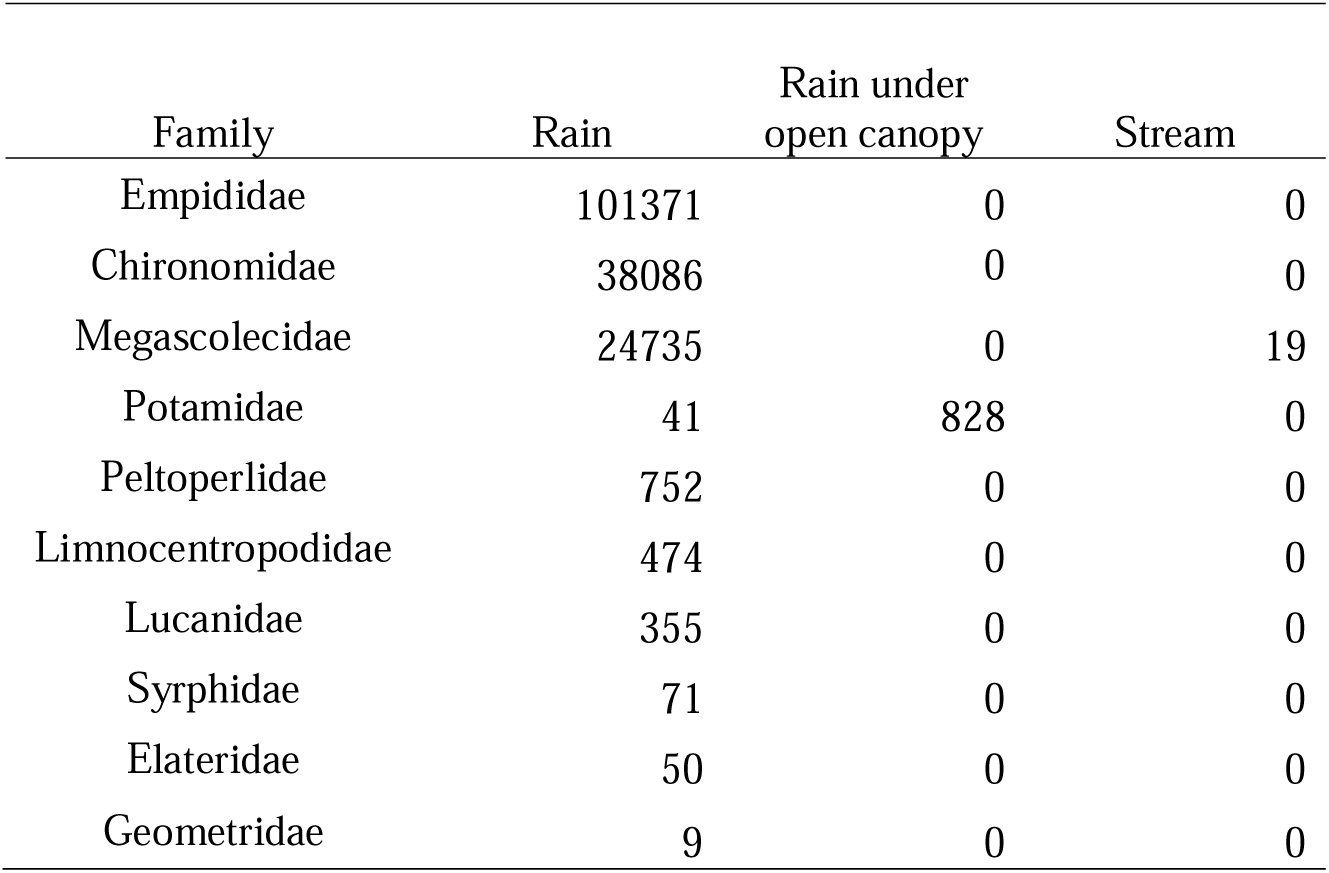
The list of detected sequence reads of the invertebrate families in the different categorical samples in Summer survey.

Figure 3 illustrates the proportion of detected families by tree species for all seasons, allowing examination of habitat preferences among taxa. Canopy drip rain samples were categorized by tree species (Fagaceae, Betulaceae, Lauraceae, Quercus, and Camellia), and the proportions of families detected from each sample type, including rain and stream samples, were plotted. Tree species at sampling locations were recorded as shown in Figure 1. Figure 3 shows that 16 out of 27 detected families were confirmed from a single sample type, with Fagaceae yielding nine families exclusively, Lauraceae one family, Betulaceae two families, and Camellia one family, while three families were detected from stream samples. Among families detected from multiple sample types, Chironomidae was detected from Betulaceae, Lauraceae, Fagaceae, and stream samples, showing the highest prevalence across sampling locations.

**Figure 3.**
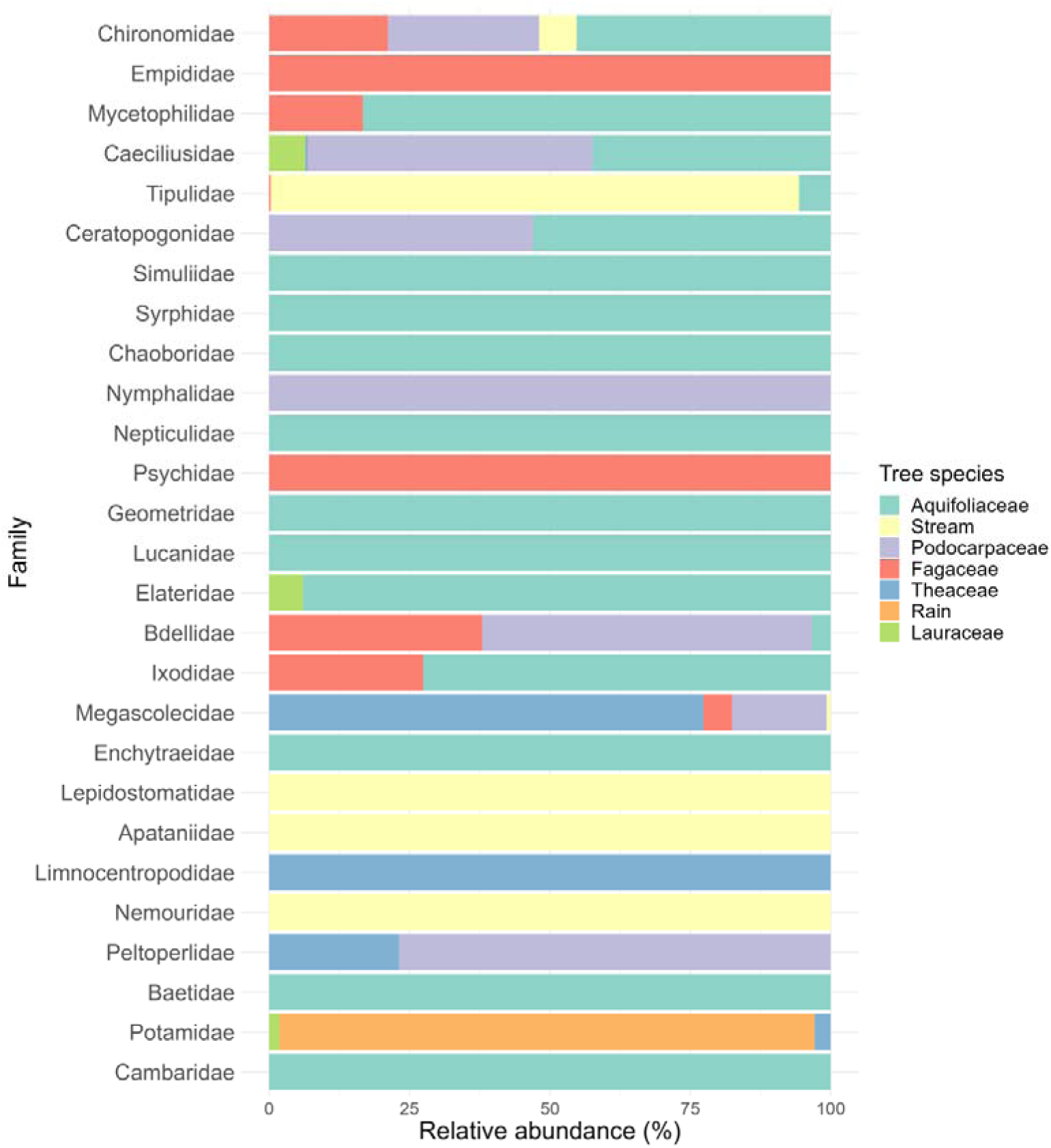
The composition of relative abundance (% reads) of the detected invertebrates in the tree species in both seasons.

Community similarity among samples is illustrated in Figure 4 using an NMDS plot, where the distance between plots indicates the degree of community similarity, with closer distances indicating higher similarity and vice versa. In Figure 4, winter survey plots are depicted as squares and summer survey plots as triangles, with summer survey plots densely clustered on the right side of the plot and winter survey plots spreading vertically but concentrated towards the left, indicating differences in community similarity between seasons, corroborated by PERMANOVA results (tree species F= 1.5678, P= 0.031, Season F=15.71, P< 0.001, interaction F=1.35, P=0.106). Additionally, regarding sampling locations, distinct differences in community similarity are observed, particularly between streams and various tree species during the winter survey.

**Figure 4.**
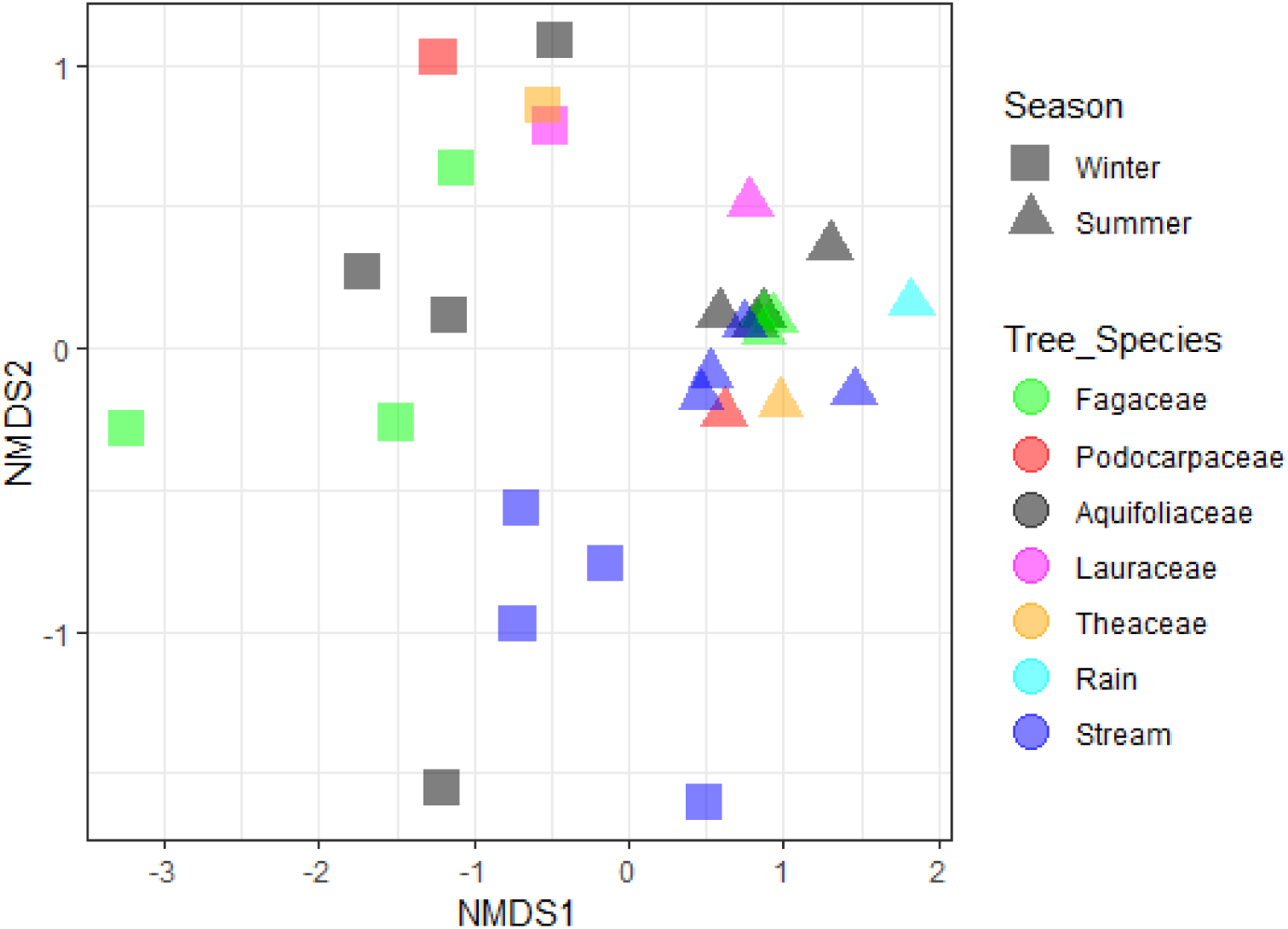
NMDS plot showing community similarity of December and June surveys (Stress value = 0.088). The different plot symbols represent sample characteristics, with tree species (including stream and rain samples) distinguished by color and sampling period (winter, summer) by shape.

### Comparison with Survey Records

The visual survey records revealed a diverse range of species, including but not limited to visually observed, sampled, and photographed species in the vicinity of the Himeji City Science Museum. By comparing the taxa at the genus and species levels detected in this study with the biological survey records of the Himeji City Science Museum, Table 4 demonstrates the comparison results. The survey records from the museum exclusively cover observations around the Himeji City Science Museum. From the sequences obtained in this study, we identified 24 genera and 11 species of biological taxa. Among these, seven genera and three species overlapped with those confirmed in the museum’s survey records. These three species were found within the seven genera. Comparing the results, approximately 29% of the genera detected in the eDNA survey matched those in the Himeji City Science Museum’s survey records, while approximately 27% of the species matched. About 70% of genera and species did not match, indicating the ability of the eDNA survey to detect species not recorded in the museum’s survey records. Furthermore, the museum’s survey records included 333 preserved species, and the overlap between the genera and species detected in our study’s two DNA surveys was approximately 3%, indicating that our study did not achieve comprehensive coverage of the museum’s survey records.

Table 4. Comparison between eDNA Survey and Himeji City Science Museum Biological Survey Records. The presence of confirmed species in the visual inspection of the Himeji City Science Museum biological survey is indicated by “〇,” while for specimens and photo surveys, the survey month is indicated. Months where the survey month coincided between the Himeji City Science Museum biological survey records and the eDNA survey are highlighted in red.

## Discussion

We found that the successful detecting DNA fragments. In the December survey, we detected 19 taxa, while in June, we detected 10 taxa. Both surveys predominantly detected aquatic invertebrates, especially families within the order Diptera. In the winter survey, rainwater passing through the canopy yielded numerous sequencing reads, while rainwater alone yielded no detections. This suggests that rainwater passing through the canopy washes away canopy-associated eDNA, and rainwater thus collected may contain this DNA, indicating the feasibility of canopy-associated eDNA surveys. Families such as Chironomidae and Libellulidae, being aquatic invertebrates, were detected from both canopy-passing rainwater and stream samples, given their larvae hatch from water. In the summer survey, we detected Lucanidae inhabiting tree trunks and decaying wood, indicating the potential detection of eDNA from substrates other than directly above the collection bucket. The occurrence of Gecarcinidae in rainwater samples suggests potential intrusion and escape of these organisms within the buckets, as no organisms were visually confirmed within the collected samples. As the buckets were placed near the Himeji City Science Museum building, it’s unlikely that eDNA from leaf litter or twigs was a significant factor (Yoneya et al. 2023). Given that Gecarcinidae are confirmed in the vicinity of the Himeji City Science Museum, the possibility arises that these organisms entered the buckets and subsequently escaped, or rain directly impacted Gecarcinidae, allowing their DNA to enter the buckets. Apart from Gecarcinidae, no sequence reads were detected in rainwater samples, predominantly detected in canopy-passing rainwater samples, akin to the winter survey. Additionally, taxa like Procambarus and Lumbricidae, typically surface-dwelling organisms, were detected in canopy-passing rainwater samples. Similar reasons to the detection of Gecarcinidae from rainwater samples are considered, indicating that although the study targeted canopy-dwelling organisms, eDNA from surface-dwelling organisms may also be captured (e.g., Ladin et al. 2021, Macher et al. 2022).

Families like Chironomidae and Lumbricidae were observed in both winter and summer surveys. Especially, Lumbricidae, prevalent in both larvae and adults in June, while only sporadic adult sightings occur in November to December. Chironomidae sightings range from March to December, suggesting detection in both winter and summer surveys. Lucanidae, found in various broadleaf forests and especially attracted to Fagaceae like Quercus and Castanopsis, were only detected in samples from Castanopsis. While this suggests Castanopsis as a preferred species for Lucanidae, it doesn’t exclude the possibility of Lucanidae being found in other tree species. Other detected taxa similarly hint at species preferences for certain tree species, with 9, 1, 2, and 1 species confirmed in Castanopsis, Machilus, Fagaceae, and Camellia, respectively. This indicates species preference concerning habitat selection across different tree species. However, the detection of multiple taxa from a single species might also suggest mere coincidence rather than inherent preferences. Additionally, factors like invertebrates species near streams/rivers being more accessible regardless of tree species and variations in rainwater angles impacting accurate representation of tree species’ canopy rainwater are plausible.

### Comparison with Visual Survey

Approximately 70% of the species detected via eDNA surveys were unrecorded in the traditional surveys, mainly comprising small-sized species like those in the order Diptera, challenging to collect. Given that eDNA surveys capture DNA left behind by organisms, irrespective of their visibility or accessibility, they complement traditional survey methods. While approximately 30% of the eDNA survey results aligned with the traditional surveys, differences in survey scope likely account for the disparity. The canopy-focused eDNA survey of this study differed from the museum’s surveys, which also included ground-dwelling species. Taxa like Coenagrionidae, found in both surveys, are observable both in trees and on the ground, hence the alignment in results. Aquatic species like *Potamon* and *Procambarus*, detectable in the museum surveys, were challenging to detect in rainwater samples, as discussed earlier. The limited alignment (only 3%) between eDNA surveys and museum records may also stem from the disparity in survey duration, with the former conducted twice compared to the latter’s multi-year span.

### Limitations and Challenges

We encountered several challenges of rainwater-utilizing eDNA analysis for canopy biological surveys. Firstly, the potential contamination of collected rainwater with forest debris and organisms, despite filtration within 24 hours, remains a concern. While visual inspection didn’t confirm organisms, the presence of debris was noted. Assessing the impact of such debris on sequence read numbers was challenging. Shortening the time between bucket setup and retrieval might mitigate this issue, but conducting operations during rainfall diminishes practicality. Secondly, varying rainfall volumes in sampling timings influenced collected water amounts, potentially affecting DNA concentrations. Investigating the relationship between collected water volume and eDNA concentration could optimize sampling methods. However, conducting such experiments amidst rainfall poses logistical challenges. Furthermore, assessing the impact of pre-survey rainfall on canopy eDNA remains challenging.

## Conclusion

In conclusion. we successfully conducted eDNA surveys on the canopy by utilizing rainwater, yet there is room to further enhance the reliability of the findings. As noted in the challenges section, conducting the operation in rainy conditions may potentially yield more reliable results. Although it may diminish practicality, reducing the time from bucket deployment to collection could mitigate external influences. Additionally, addressing rainfall prior to survey days presents challenges due to the difficulty in adjusting the period between rainfalls. To tackle this, conducting multiple surveys and recording precipitation before survey days can help understand this pattern. Narrowing the study scope to a single leaf within the forest could evaluate the impact of pre-survey rainfall. By injecting water onto leaves, collecting the water that passes through, and analyzing eDNA concentration, it’s possible to assess the extent of eDNA adhering to forest leaves. This approach, conducted at predetermined intervals over several hours and days, could determine the level of eDNA deposition on forest leaves, providing crucial insights for optimizing eDNA surveys using rainwater and enhancing result reliability. Given the substantial costs associated with traditional canopy biodiversity surveys in large forests like tropical rainforests, this methodology offers a relatively straightforward approach, potentially capturing previously unidentified species.

## Data Availability Statement

This raw data was deposited in DRA: DRA**.

## Author contributions

TM and HD designed the study; TM, NM, and CN collected the data; TM, CN and HD analyzed the data; TM and HD interpreted the results; TM, NM and HD wrote the manuscript.

## Acknowledgments

We thank the Himeji City Science Museum in Himeji City, Hyogo Prefecture for providing the sampling location and observation data.

